# Optimizing linear ion trap data independent acquisition towards single cell proteomics

**DOI:** 10.1101/2023.02.21.529444

**Authors:** Teeradon Phlairaharn, Zilu Ye, Elena Krismer, Anna-Kathrine Pedersen, Maik Pietzner, Jesper V. Olsen, Erwin M. Schoof, Brian C. Searle

**Affiliations:** The Novo Nordisk Foundation Center for Protein Research, Faculty of Health Sciences, University of Copenhagen, Copenhagen, Denmark; Technical University of Munich, Germany; TUM School of Natural Sciences, Department of Bioscience; Computational Medicine, Berlin Institute of Health at Charité–Universitätsmedizin Berlin, Berlin, Germany; Department of Biotechnology and Biomedicine, Technical University of Denmark, Lyngby, Denmark; Pelotonia Institute for Immuno-Oncology, The Ohio State University Comprehensive Cancer Center, Columbus, Ohio 43210, United States; Department of Biomedical Informatics, The Ohio State University, Columbus, Ohio 43210, United States

## Abstract

A linear ion trap (LIT) is an affordable, robust mass spectrometer that proves fast scanning speed and high sensitivity, where its primary disadvantage is inferior mass accuracy compared to more commonly used time-of-flight (TOF) or orbitrap (OT) mass analyzers. Previous efforts to utilize the LIT for low-input proteomics analysis still rely on either built-in OTs for collecting precursor data or OT-based library generation. Here, we demonstrate the potential versatility of the LIT for low-input proteomics as a stand-alone mass analyzer for all mass spectrometry measurements, including library generation. To test this approach, we first optimized LIT data acquisition methods and performed library-free searches with and without entrapment peptides to evaluate both the detection and quantification accuracy. We then generated matrix-matched calibration curves to estimate the lower limit of quantification using only 10 ng of starting material. While LIT-MS1 measurements provided poor quantitative accuracy, LIT-MS2 measurements were quantitatively accurate down to 0.5 ng on column. Finally, we optimized a suitable strategy for spectral library generation from low-input material, which we used to analyze single-cell samples by LIT-DIA using LIT-based libraries generated from as few as 40 cells.

## INTRODUCTION

The latest advances in nanoflow liquid chromatography-tandem mass spectrometry (LC-MS/MS)^1,2^ have brought low-input proteomics into the spotlight. Inherent optimization of upstream methods such as sample preparation,^3–5^ chromatography,^6^ and data acquisition strategies^7–10^ is fundamental for obtaining the deepest proteome coverage while maintaining high quantitative accuracy. To achieve high sensitivity and robustness essential in MS-based single-cell proteomics (SCP-MS),^11,12^ Budnik et al^7^ have shown the potential of using a multiplexing approach in SCP-MS with data-dependent acquisition (DDA).^8^ However, quantitative precision can be limited when using tandem mass tags (TMT) with a carrier proteome in multiplexing experiments.^13,14^

In recent years, data-independent acquisition (DIA) has become increasingly popular for quantitative proteomics where co-eluting peptides are co-analyzed with wider isolation windows and fragmented together regardless of their precursor intensities.^15,16^ For this reason, DIA is an unbiased mass spectrometric approach that produces fewer missing values than DDA, improving the reproducibility of MS-based proteomics experiments. Most orbitrap (OT)-based DIA methods for low-input material are implemented with very wide precursor isolation windows (up to 60 m/z)^10,17^ to ensure a high enough ion injection time to measure low abundant ions while maintaining an overall acquisition scan cycle time generating sufficient measurement points across a chromatographic peak. However, the tradeoff of using expansive isolation windows is the loss in precursor selectivity producing highly chimeric MS2 spectra.

Although DIA is typically performed on high-resolution mass spectrometers, historically the method was first implemented on linear ion traps (LIT).^18^ Compared to the OT, the LIT has proven to be a more sensitive and faster scanning mass analyzer.^19,20^ While the limited mass accuracy of LITs cannot make use of neutron-encoded isobaric tagging quantitative methods,^21^ LITs have sufficient resolution to quantify peptides using iTRAQ or 6-plex TMT in MS3 measurements,^22^ which Park et al demonstrated can be applied to single cells.^23^ Additionally, LITs are capable of accurately quantifying fragment ions with low-input proteomics experiments using both parallel reaction monitoring (PRM)^24^ and DIA.^25,26^ However, all of these LIT-based low-input proteomics methods require either precursor ions to still be collected using a built-in OT or a spectrum library to be collected using an OT.

There is a growing interest in library-free DIA analysis,^27^ which relies on linking co-eluting precursor and fragment ions: a task made easier by high-resolution mass spectrometry. On the other hand, spectrum libraries enable peptide-centric analysis^28^ and may improve both sensitivity and specificity with low-resolution LIT analyzers. Offline fractionation^29–32^ (OF) by peptide chromatographic methods that are orthogonal to the online low pH reversed-phase chromatography is the optional upstream method to reduce sample complexity and generate high-quality sample-specific spectral libraries^33^ to significantly enhance proteome coverage in proteomics studies. However, drawbacks of extensive OF are the additional effort, instrument time required, and especially the starting material requirements, which renders OF incompatible with limited-input experiments such as Laser Capture Micro-dissected tissue samples,^34–36^ flow-sorted sub-populations of cell types^37^ or analysis of rare single cells, where material available for OF is typically insufficient.

In this work, we demonstrate a sensitive method to generate moderately deep offline high-pH reversed-phase fractionation (OHPF) libraries for LIT-based DIA analysis from as little as 50 ng of starting material. Besides OHPF, we use gas-phase fractionation (GPF)^38–40^ to generate a chromatographically-matched DIA-based library using the LIT. While OHPF is incompatible with sample loads < 50ng, we show that GPF libraries for DIA analysis can be generated from as little as 4 ng of pooled peptide material per injection. We further prove the sensitivity of this approach using input material from as little as 10 cells to mimic the peptide fragmentation spectra^41^ from SCP-MS. Finally, we demonstrate the robustness, reproducibility, and sensitivity of this approach by analyzing single cells using only a LIT for both MS1 and MS2, detecting more than 950 peptides in 31 minutes with a LIT DIA-based chromatogram library from the equivalent of 16 ng starting material. These results indicate the potential viability of using inexpensive LIT-only instruments for single-cell DIA-based mass spectrometry.

## EXPERIMENTAL

### Sample Preparation

For the dilution series of HeLa protein lysate, we used Pierce HeLa Protein Digest Standard (Thermo Scientific, Catalog number: 88328). For the yeast protein digest in the mixed proteome experiment, MS-compatible yeast intact (undigested) extracts were purchased from Promega (Catalog number: V7341) and processed according to the technical manual.

HEL 92.1.7 (ATCC TIB-180) cells were cultured in Gibco RPMI 1640 with GlutaMAX supplement (Thermo Scientific) containing 10% fetal bovine serum (FBS, Thermo Scientific) and 100 µg ml^−1^ Penicillin–Streptomycin (Thermo Scientific). Cells were kept at 37 °C in a humidified incubator with 5% CO2. For cell culture maintenance cultures were maintained at 0.5 × 10^6^ with medium renewal and transferred into a new T75 flask every 2-3 days. For single-cell and bulk experiments samples prepared with the cellenONE platform, the cell suspension was spun down, and followed by removing the supernatant. The cell pellet was washed twice by adding 10 mL of DPBS and spindown at 300 xg for 5 minutes at 4°C. To prepare the cell suspension for cell sorting using cellenONE, the cell pellet after the washing step was resuspended in DPBS at an appropriate density. A 96 wells plate was preloaded with 2 µL n-Hexadecane (Thermo Scientific). 300 nl lysis buffer containing 0.1% DDM, 5 mM TCEP in 100 mM TEAB, 20 mM CAA in TEAB, 100 ng/nl trypsin, and 100 ng/nl Lys-C was preloaded into 96 wells plate. HEL 92.1.7 cells were sorted into the 96 wells plate and incubated at 25 °C overnight. After the incubation, 8 µL of 0.1% TFA was added to acidify each sample. Lastly, the 96 wells plate was incubated at 4 °C to solidify the n-Hexadecane.

EvoTip PURE loading tips were activated by adding 25 µL buffer B, centrifugation at 600 xg, and conditioning in 1-propanol for 5 minutes. Then, 30 µL buffer A was added to each EvoTip, followed by centrifugation at 650 xg for 1 min. Each sample from the 96 wells plate with solidified n-Hexadecane was then loaded into an EvoTip, followed by centrifugation at 650 xg for 1 min and then two additional centrifugation steps at 650 xg for 1 min after adding 30 µL buffer A. Lastly, 150 µL buffer A was added to each EvoTip followed by a final centrifugation step for 5 sec at 650 xg.

### Off-line moderately high-pH reversed-phase fractionation (OHPF) and library generation

To generate a spectral library for this study, we fractionated 50 ng, 100 ng, and 1000 ng of HeLa protein digest with an OF into 8 fractions on 100 minutes gradient at a 1.5 µL/min flow rate. As the fractionator, we connected the Ultimate 3000 HPLC (Dionex, Sunnyvale, CA, USA) with a 150 mm EVO C18 column with a particle size of 2.6 µm and an inner diameter of 0.30 mm. We increased the concentration of buffer B (10 mM TEAB in 80% ACN at pH 8.5) from 3% to 40% in 57 minutes, 60% in 5 minutes, and 95% in 10 minutes. The concentration of buffer B was constant at 95% for 10 minutes and steadily decreased to 3% in 10 minutes. The fractions were loaded into the EvoTip PURE loading tips and acquired in DDA with a 31-minute stepped pre-formed gradient (Whisper 100 40 SPD) for spectral libraries generation. For DDA-based spectral libraries, raw data from the OF were used to generate a library via the Pulsar search engine in Spectronaut^42^.

### Gas-phase fractionation (GPF-DIA) and spectral library generation

To generate the chromatogram library for this study, we applied the gas-phase fractionation approach to build DIA-based chromatogram libraries from the total 40 ng of HeLa peptide digest and equivalent to 10 (40 cells), and 16 (64 cells) ng of HEL peptide digest. Briefly, the same amount of peptides digest was injected 4 times covering 100 m/z for every peptide between 500 - 900 m/z, and acquired with a 31-minutes stepped pre-formed gradient (Whisper 100 40 SPD) and these measurements were combined into a single DIA-based library. For chromatogram library generation, raw data from GPF experiments were searched via Spectronaut.

### LC-MS/MS Analysis

We used an Evosep ONE system for this study and analyzed single-cell tryptic digest and diluted HeLa peptide digest with a 31-minutes stepped pre-formed gradient (Whisper 100 40 SPD) eluting peptides at a 100 nl/min flowrate. We used a 15 cm x 75 µm ID column (PepSep) with 1.5 µm C18 beads (Dr. Maisch, Germany) and a 10 μm ID electrospray emitter (PepSep). Mobile phases A and B were 0.1 % formic acid in water and 0.1 % in ACN. The LC system was coupled online to an orbitrap Eclipse Tribrid Mass Spectrometer (ThermoFisher Scientific) via an EasySpray ion source connected to a FAIMSPro device. We set the nanospray ionization voltage to 2300 volts, the FAIMSPro gas flow to static (3.6 L/min), and the temperature of the ion transfer tube to 240 °C. We applied the FAIMSPro ion mobility device’s standard resolution and used a compensation voltage (CV) of −45 volts. NCE was configured to 33. Specific configurations for the instrument acquisition settings are below.

#### DIA

The scan sequence began with an MS1 spectrum on LIT (Scan range 400–1000 Th, automatic gain control (AGC) target of 300%, maximum injection time 10 ms, RF lens 40%). The precursor’s mass range for MS2 analysis on LIT was set from 500 to 900 Th and the scan range from 200 - 1200 Th. MS2 analysis consisted of higher-energy collisional dissociation (HCD), and MS2 AGC was set to standard. the isolation window was set based on scanning speeds on the MS2 level. (Supporting information: Table S2).

#### GPF-DIA

The scan sequence began with an MS1 spectrum on LIT (Scan range 400–1000 Th, automatic gain control (AGC) target of 300%, maximum injection time 10 ms, RF lens 40%). The precursor’s mass range for MS2 analysis on LIT covered 100 m/z for every peptide between 500 - 900 m/z (500 - 600 m/z, 600 - 700 m/z, 700 - 800 m/z, and 800 - 900 m/z), and the scan range from 200 - 1200 Th. MS2 analysis consisted of higher-energy collisional dissociation (HCD), and MS2 AGC was set to standard. the isolation window was set based on scanning speeds on the MS2 level. (Supporting information: Table S2).

#### DDA

The scan sequence began with an MS1 spectrum on OT (120.000 resolution, Scan range 375–1500 Th, automatic gain control (AGC) target of 100%, maximum injection time 50 ms, RF lens 30%). Precursors for MS2 were selected in a Top10 fashion, and only charge states 2-6 were included. Precursors were isolated for MS2 analysis using 1.4m/z isolation windows, fragmented using a normalized collision energy of 30 (HCD), and MS2 AGC was set to 300%, with a maximum injection time of 35ms. Scan range was fixed from 200-1400, with LIT scan speed set to ‘*Normal*’.

The supporting information for additionally tested methods in Supplementary Table S1 is listed as Supplementary Methods.

### Data analysis

#### Raw LC-MS/MS data analysis and downstream proteomics data analysis

For the DDA analysis of HeLa protein lysate global quality control and determination of separation efficiency of our OHPF system, we used MaxQuant^43^ (version 2.1.3.0). We used Spectronaut ^42^ (version 17.0.221020.50606) (Biognosys)to analyze DIA raw data in directDIA mode (Library free) and library-based searches, with additional quantification performed using EncyclopeDIA^44^ (version 2.12.30). False-discovery rates (FDRs) on peptide spectral match (PSM), peptide, and protein levels were equal to 1%. Trypsin/P was selected for enzyme specificity. The allowed peptide length in the search space was between seven and fifty-two. A maximum of two missed cleavages were allowed. Cysteine carbamidomethyl was selected as a fixed modification and for the variable modifications, N-terminal acetylation and methionine oxidation were selected. For other parameters, we applied the default settings. EncyclopeDIA was configured to read Spectronaut search results and perform re-integration, transition refinement, and quantification based on raw data. To this end, EncyclopeDIA used precursor, fragment, and library mass tolerances of 0.4 AMU, and configured to search within a 1-minute window around Spectronaut detections with additional FDR assessment disabled. Output tables from Spectronaut and EncyclopeDIA (for DIA analysis) and Maxquant (DDA analysis for separation efficiency) were imported into RStudio (Version 2022.07.2) for downstream analysis and visualization.

## RESULTS AND DISCUSSION

The main goal of this study was to evaluate using standalone LIT mass analyzers for low-input and single-cell proteomics from an analytical perspective. While limited in mass accuracy, LITs are both less expensive and easier to maintain than high-resolution orbitrap or time-of-flight mass spectrometers. These criteria make a standalone LIT instrument a potentially ideal platform for high-throughput single-cell work where single experiments will require thousands of injections. Towards this end, we assessed several parameters: 1) we determined optimal DIA methods for operating the LIT analyzer, 2) we tested if search engines designed for high-resolution DIA data could accurately estimate false discovery rates, and 3) we estimated global lower limits of quantitation for LIT measurements in low-input scenarios.

### Assessing optimal windowing and estimating confidence in FDR assessments

The Thermo LIT analyzer in the Eclipse has five different scanning modes: turbo, rapid, normal, enhanced, and zoom, in order of scan speed. We analyzed 1 ng and 10 ng samples of HeLa protein lysate using LIT on Whisper 100 40 SPD,^45^ where we varied the precursor isolation window width (and thus the total number of windows), while keeping the cycle time (~2 seconds) and overall precursor mass range (500-900 m/z) the same. Analyzing these with Spectronaut in directDIA mode, we found that detections were maximized using different scanning modes for each sample concentration where 10 ng samples produced the greatest number of peptide detections using 80× 5 m/z turbo scans per cycle. In contrast, the 1 ng sample detections were maximized using 40× 10 m/z turbo scans per cycle (**Figure 1**).

**Figure 1.**
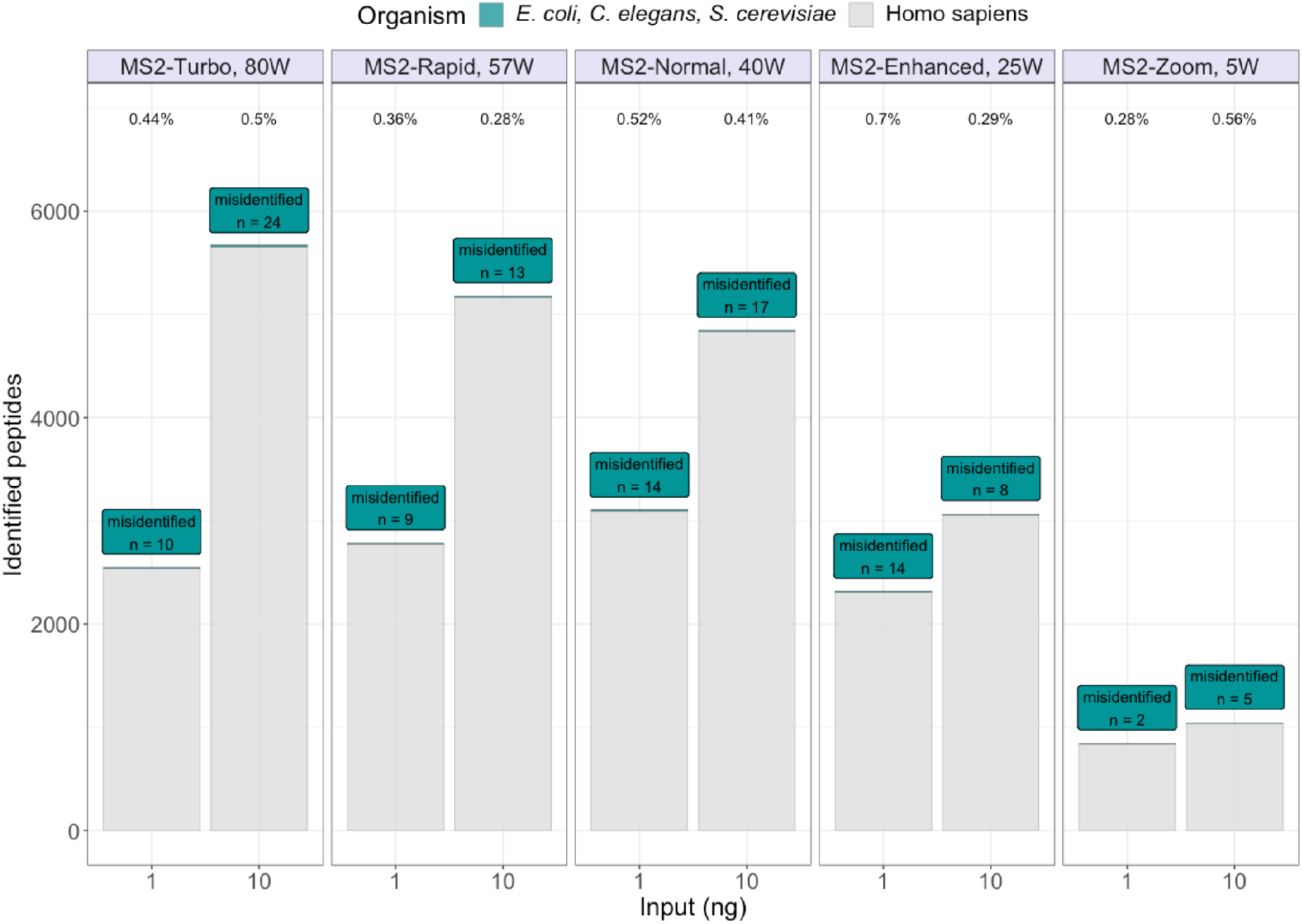
FDR confidence estimation using a LIT for low-input experiments. The number of detected peptides when searching LIT-DIA datasets from 1 ng and 10 ng protein on column with different scan settings with windows that are matched to maintain approximately a 2-second cycle time. Searches were performed using the entrapment approach, where in addition to searching a *H. sapiens* database, peptides from *E. coli, C. elegans*, and *S. cerevisiae* were also considered. The number of misidentified entrapment peptides, as well as the ratio of entrapment to total detected peptides are shown above bars.

Due to the low mass accuracy of LITs, we were concerned that Spectronaut, designed to analyze high-resolution DIA data, might incorrectly assess the peptide false discovery rate in our experiments. To test this, we performed an entrapment experiment^46–51^ to measure the accuracy of target/decoy-based false discovery rate (FDR) estimations^52^ these experiments. We configured Spectronaut to filter at a target/decoy estimated FDR of 1% at the precursor and protein levels, and a human protein sequence database (5,142,722 tryptic peptides) using additional entrapment sequences (4,463,824 total entrapment tryptic peptides) from Escherichia coli (*E. coli)*, Caenorhabditis elegans *(C. elegans) and* Saccharomyces cerevisiae *(S. cerevisiae)*. Since the samples should only contain human peptides, assignments made to entrapment peptides indicate false positives. The ratio of entrapment to total peptides is 46.4%, which closely matches to the 1% of the average assignments to entrapment peptides (0.434%) shown in **Figure 1**.

After validating that the FDR estimates were reasonably correct across all scan types, we next sought to determine the optimal settings for LIT measurements using 1, 10, and 100 ng of input material. To do this, we performed several experiments varying both MS1 and MS2 scan modes for each input level, keeping the overall cycle time between 1.9 and 2.8 seconds to achieve sufficient points across the peak for quantification (**Supplementary Table S1**). These results suggested that for a 1 ng injection, the ‘*Enhanced*’ scanning mode on MS1 and ‘*Normal*’ scanning mode on MS2 is optimal. For 10 ng, ‘*Normal*’ on MS1 and ‘*Rapid*’ on MS2 performed best, while for 100 ng, ‘*Rapid*’ on MS1 and ‘*Rapid*’ on MS2 scans produced the best results (**Table 1**).

**Table 1.** Optimal parameters for LIT-DIA for different sample loads that maintain approximately 2-second cycle times.

| Input | MS1 scanning mode | MS1 reached max fill time | MS2 scanning mode | MS2 reached max fill time | Window size | Cycle time |
| --- | --- | --- | --- | --- | --- | --- |
| 1 ng | *Enhanced* | 62.4 % | *Normal* | 99.9 % | 10 m/z | 2.08 s |
| 10 ng | *Normal* | 30.9 % | *Rapid* | 99.9 % | 7 m/z | 1.98 s |
| 100 ng | *Rapid* | 21 % | *Rapid* | 88.3 % | 7 m/z | 1.94 s |

### Assessing the quantitative accuracy of low-input LIT-DIA measurements

Compared to our previous results using OT for MS1 and LIT for MS2,^26^ the detected peptides analyzed by LIT alone expected are somewhat lower (**Figure 2A**). However, without the OT MS1 scan, the cycle time is sufficiently fast to achieve at least 8 points per peak (PPP) needed for accurate fragment-level quantification.^53^ We then evaluated the quantitative reproducibility across replicates by calculating the Pearson correlation coefficient (**Figure 2B**). From this, we found that randomly paired replicates produced very similar quantitative results, even with only 1 ng of peptides on column. We found that both precursor and protein-level quantification correlated well with the amount of sample on column (**Figure 2C**), where the intensities for MS1 were typically higher and fell within a narrower range than MS2 intensities.

**Figure 2.**
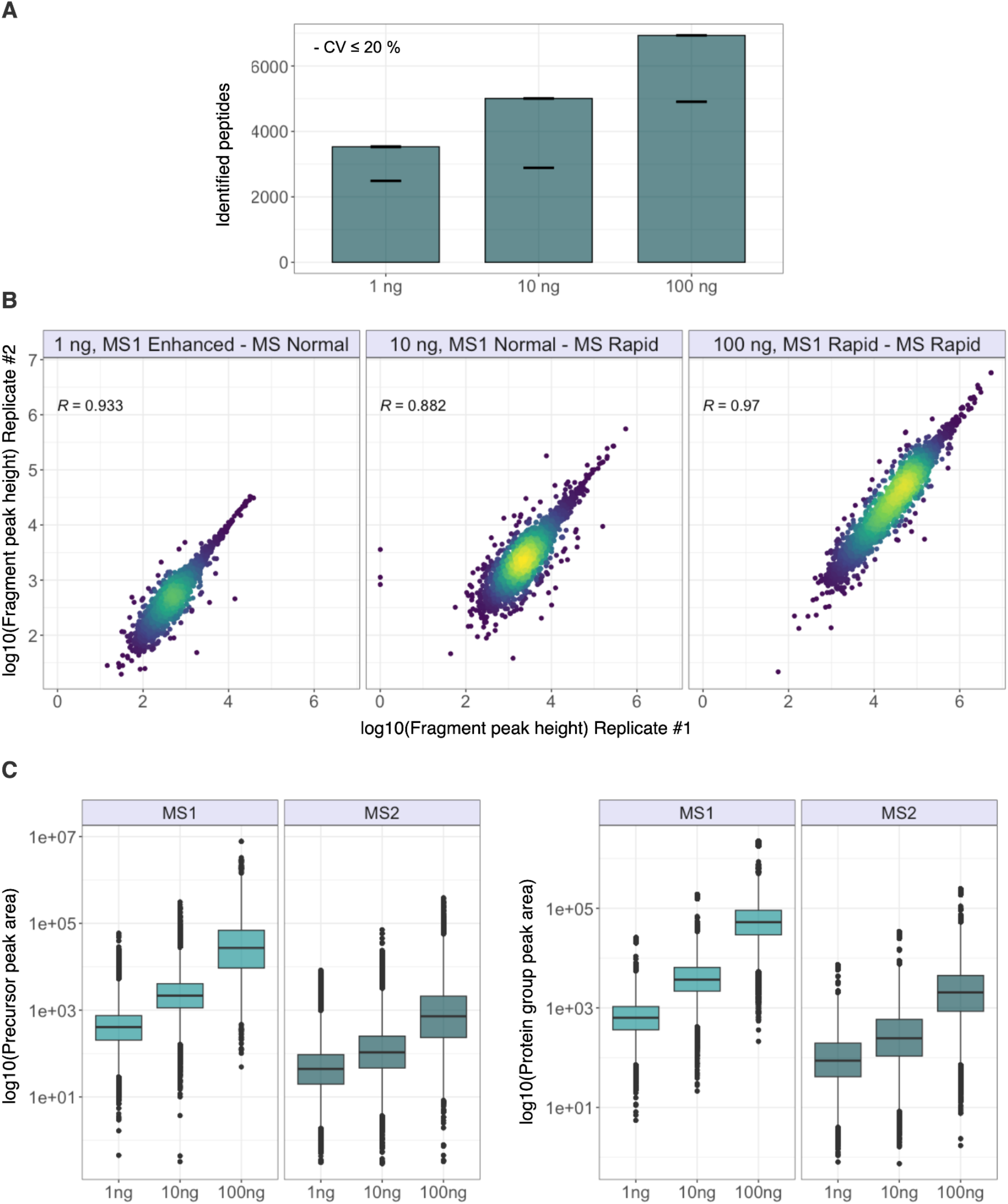
Repeatability of LIT-DIA measurements at different loads. (**A**) The number of detected peptides from triplicate 1, 10, and 100 ng of HeLa protein lysate. Error bars at the height of the bars indicate the spread of values in the triplicate, while error bars below the bars indicate the spread of high-confidence peptide detections with intensity coefficient of variances below 20%. (**B**) The scatter in individual fragment ion peak height reproducibility between two replicates for the same lysate samples. (**C**) The overall distribution of summed peak areas at either the precursor or protein levels.

Consistency with dilution in the solvent can assess sensitivity but not quantitative accuracy. As target peptides decrease in signal with dilution, the background also decreases by an equal amount, which reduces the effect of interference. To test quantitative accuracy, we instead diluted an *S. cerevisiae* protein digest into a HeLa protein digest similar to a matched matrix calibration curve.^54^ Maintaining a total of 10 ng on column, we measured mixtures of yeast and HeLa protein digest in eight ratios (100%:0%, 50%:50%, 20%:80%, 10%:90%, 5%:95%, 2%:98%, 1%:99%, and 0%:100%). From the 10 ng of yeast protein digest, we detected 551 proteins using LIT-DIA, where 432 proteins had a coefficient of variation (CV) < 20% across all replicates. While we created the dilution series based on total protein mass, the relative amount of signal observed by the mass spectrometer must be normalized because the yeast and human proteomes are of differing complexities. For this, we reported quantification relative to the average MS1 (from Spectronaut) and MS2 (from EncyclopeDIA) ratio of the 50%:50% sample measurement to the 100% yeast sample.

As shown in **Figure 3**, we observed poor quantitative performance with MS1 measurements, most likely due to low precursor mass accuracy. However, MS2-based quantification was significantly better, even when analyzing as little as 0.5 ng of input material. This improved quantification is likely due to the increased selectivity from windowing by isolating precursors from a narrow m/z range, which appears to overcome the lower LIT mass accuracy. Interestingly, with Spectronaut, low-quality signals seemed to introduce higher degrees of noise into the quantitative signal, most likely due to the addition of interference from the human background proteome. We observed this in both MS1 and MS2 measurements, where the quantities assigned to yeast proteins appeared to increase at lower yeast:human ratios. To test this, we used EncyclopeDIA to re-quantify peptide detections made by Spectronaut. While EncyclopeDIA does not currently support LIT-DIA for searching, we used EncyclopeDIA to perform MS2 quantitative signal extraction and transition refinement at the retention time locations for detections made by Spectronaut. Interestingly, these results were both more precise around the expected ratios, as well as having better accuracy for peptides below 0.5 ng on column. This suggests that a hybrid Spectronaut/EncyclopeDIA search and quantification workflow may improve overall results with LIT-DIA.

**Figure 3.**
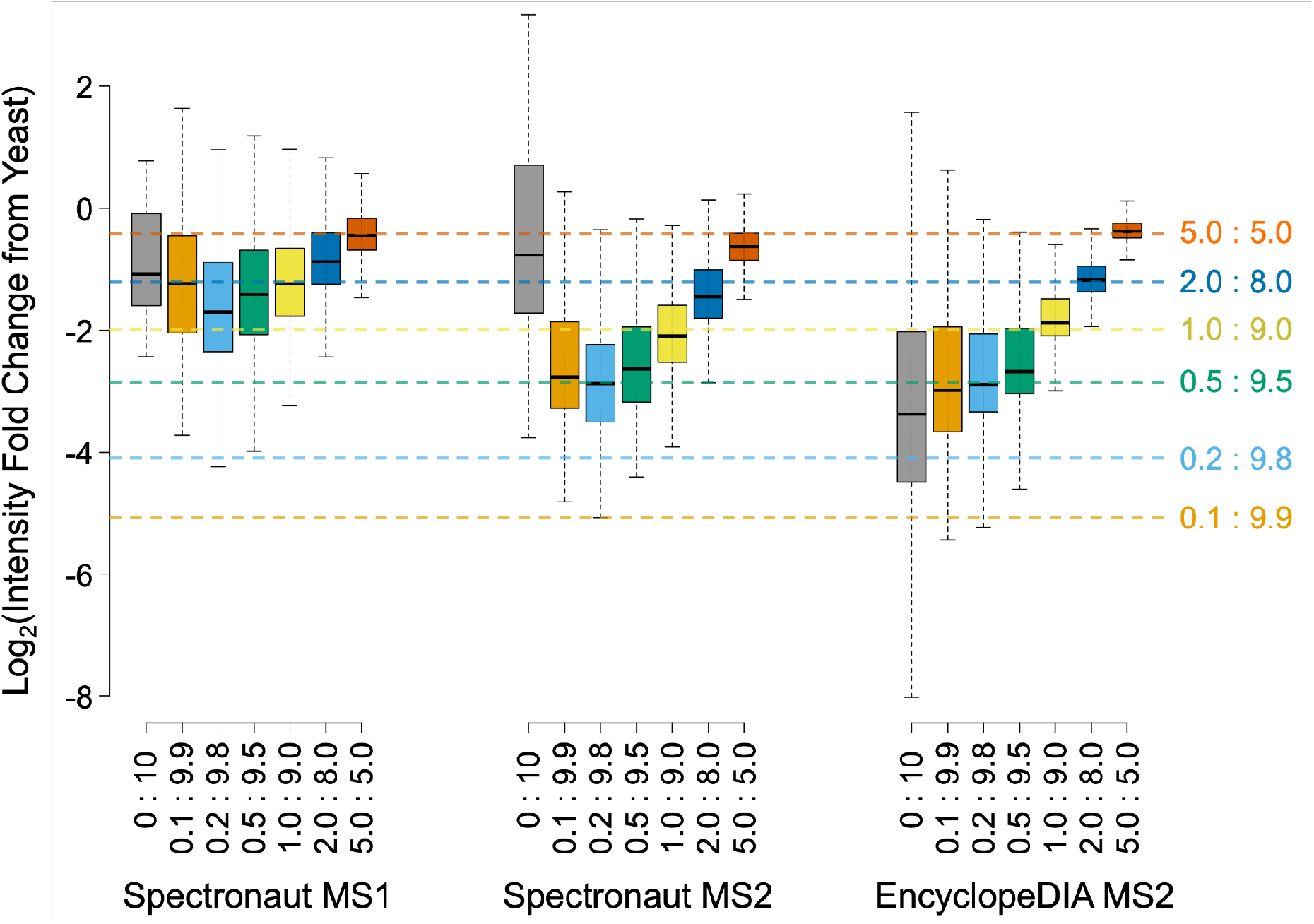
Quantitative MS1 and MS2 accuracy in low-input matrix-matched proteomes with LIT-DIA. Boxplots show the spread of diluted yeast protein intensities relative to 100% yeast in a HeLa background where boxes indicate medians and interquartile ranges and outliers are removed for clarity. Expected ratios after normalization are indicated as dashed colored lines.

### Improvement in the number of detected peptides via spectral library-based LIT-DIA

While spectrum libraries are known to improve peptide detection rates with DIA, developing these libraries with limited sample material can add additional challenges. Offline high-pH reversed-phase peptide fractionation is a standard and feasible method to generate a spectral library from both DDA and DIA data. However, a lossless and compact system is required to recover low-input materials for high-sensitivity analysis, which is generally difficult to achieve. Alternatively, a chromatogram library-based strategy using GPF-DIA is potentially suitable for this type of experiment because samples can be directly injected into the mass spectrometer. The chromatogram library approach may be an effective alternative strategy to analyze limited input proteomics-based libraries because it requires less material for library generation.

We tested several different approaches to generating spectrum libraries using different fractionation strategies: either OHPF to generate DDA-based spectral libraries from 50, 100, and 1000 ng input material, or GPF-DIA-based chromatogram libraries from 40 ng input materials (4x fractions each covering 100 m/z isolation windows between 500-900 m/z). Analyzing single-shot HeLa runs with LIT-DIA using different input amounts, we found that the number of detected peptides scaled with library input, where OHPF libraries were superior to the GPF-based library at higher loads (10 ng and 100 ng) but comparable at the 1 ng load (**Figure 4A**).

**Figure 4.**
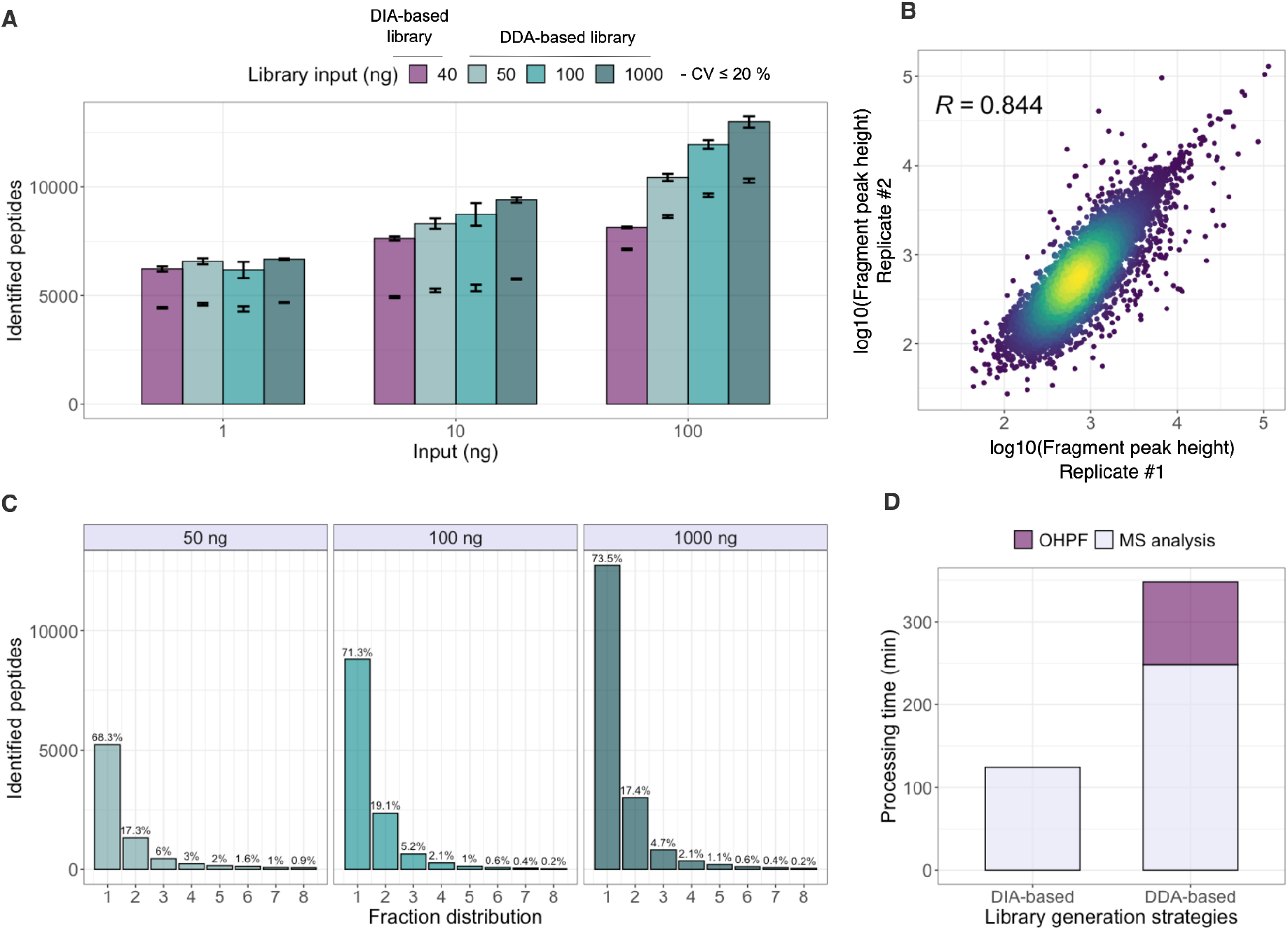
The effect of library generation approach on LIT-DIA dataset characteristics. **(A)** The number of detections from 1, 10, and 100 ng of HeLa protein lysate injections based on library generation approach. The libraries were generated by either GPF LIT-DIA using 40 ng of starting material, or OHPF LIT-DDA using 50, 100, or 1000 ng of starting material. Error bars at the height of the bars indicate the spread of values in the triplicate, while error bars below the bars indicate the spread of high-confidence peptide detections with intensity coefficient of variances below 20%. (**B**) The scatter in individual fragment ion peak height reproducibility between two replicate LIT-DDA OHPF libraries from 50 ng of starting material. (**C**) The count of detected peptides sorted by the total number of fractions required to assign at least 75% of the total peptide intensity. (**D**) The total amount of time required to generate DIA- and DDA-based libraries, including fractionation and mass spectrometry analysis.

To determine the reproducibility of our OHPF, we performed OHPF from 50 ng input material into 8 fractions in duplicate and analyzed them individually with LIT-DIA in directDIA mode. We found that the technical reproducibility of the OHPF was robust with Pearson’s *R* = 0.844 (**Figure 4B**). Due to the inevitable sample loss during offline fractionation, we tested the GPF-DIA approach to generate a chromatogram library for limited material studies. The main advantage of generating a chromatogram library using the GPF-DIA approach is that the representative samples can be injected directly into the mass spectrometer, omitting the need for an offline fractionation system. Importantly, as there are no sample losses during the procedure: it only requires minimal sample amounts to build a chromatogram library and boost the number of detected peptides using a spectral library-based approach.^55^

The aim of performing OHPF for library generation is to stretch the complexity of biological samples into different fractions, making it easier to detect low-abundance peptides. In this study, we used a moderately high-pH buffer B (1% TEAB in 80% ACN at pH 8.5) to perform offline high-pH reversed-phase fractionation instead of a more classical high-pH buffer at pH 10 as described previously.^31,32^ As most commercially available HPLC columns are silica-based, high-pH mobile phase leads to column degradation^56^ affecting the reproducibility of analysis and consequently decreasing the efficiency of orthogonal separation for reversed-phase liquid chromatography.^57^ We determined the separation efficiency of our OHPF setup as described by Kulak and Geyer et al.^32^ Here, the percentage of peptides that have at least 75% of their total intensity identified in a single fraction was 68.3% from 50 ng starting material, 71.3% from 100 ng starting material, and 73.5% from 1000 ng starting material (**Figure 4C**). A final consideration when constructing spectrum libraries is library generation time. For OHPF-based libraries, we separated the sample into 8 concatenated fractions using a 100 minutes offline LC gradient, which thusly requires eight LC-MS/MS runs to generate the spectral library. For the GPF-based chromatogram library, we only need to inject each sample 4 times (once for each m/z range), thereby cutting instrument time in half (**Figure 4D**).

### Improvement of SCP-MS on LIT-DIA by searching against chromatogram library

We evaluated the effect of library generation with single-cell proteomes and 10-cell pools of the HEL human erythroblast cell line using a standardized sample preparation workflow based on cell sorting in microliter scale with the cellenONE platform in a 96 well-plate format, enabling label-free single-cell proteomics sample preparation.^58^ For this analysis, we configured the LIT to acquire data with ‘*Normal*’ on MS1 scanning mode and ‘*Rapid*’ on MS2 scanning mode. From the single-cell analysis using LIT-DIA in directDIA mode, we detected more than 400 peptides. By searching against a DIA-based chromatogram library created from 64 cells as a starting material, we were able to increase the number of detected peptides to over 1,100 (**Figure 5A**). Interestingly, in the bulk analysis of 10 HEL cells, there is sufficient material for the directDIA mode to outperform searching against the low-input GFP-based library (**Figure 5B**). However, the higher input libraries almost doubled the number of peptide detections versus directDIA. Our findings confirm previous reports^59^ that scaling the input sample for DDA-based spectral libraries or DIA-based chromatogram libraries based on the expected load can improve the number of detected peptides in low-input proteomics analysis.

**Figure 5.**
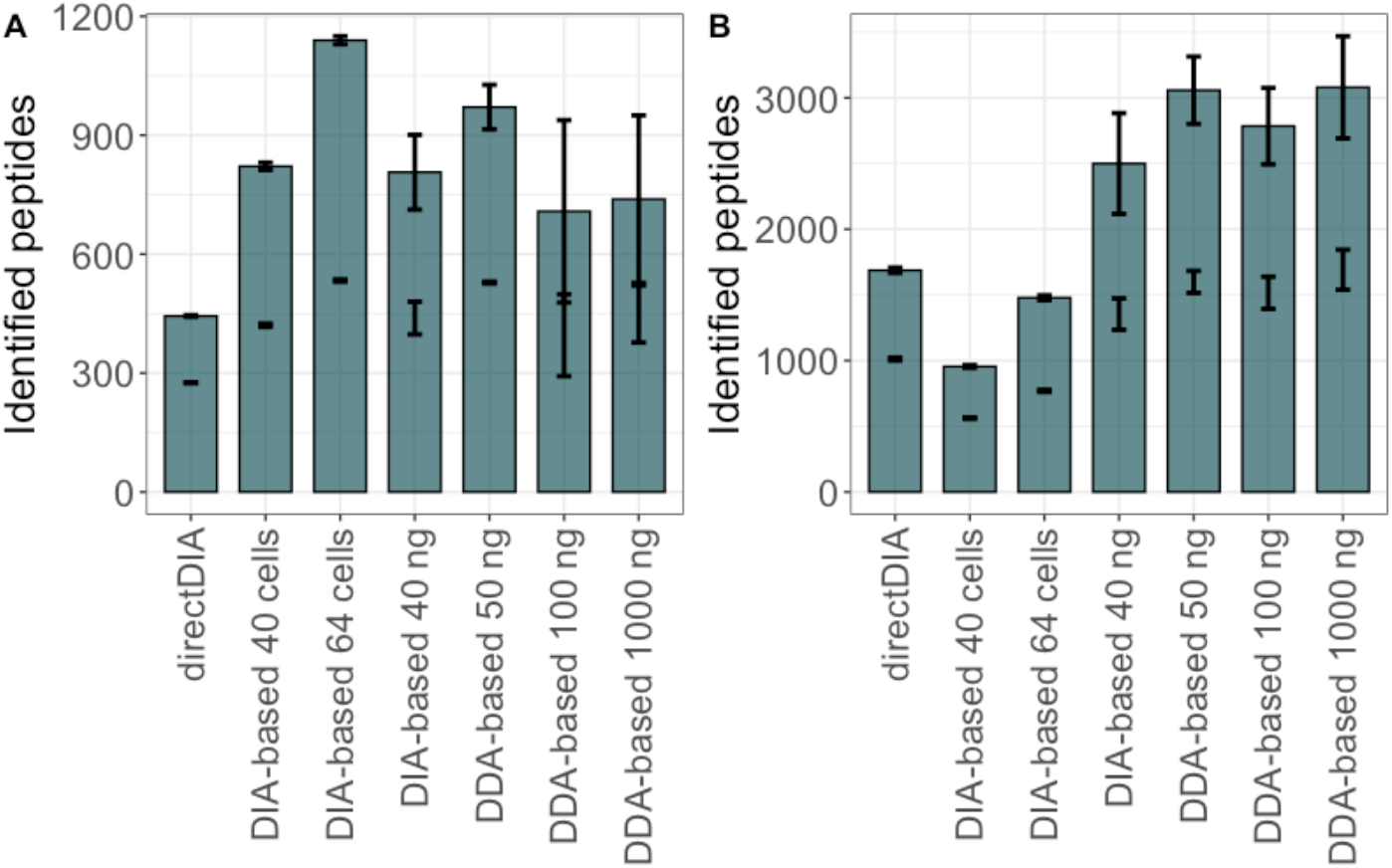
The number of peptides detected in triplicate (A) single-cell injections and (B) 10-cell pools searched using different library generation strategies. Cells were sorted via cellenONE and analyzed by LIT-DIA on Whisper 100 40 SPD and searched in directDIA mode, or searched against DDA-based spectral libraries and DIA-based chromatogram libraries generated from different input levels. Error bars at the height of the bars indicate the spread of values in the triplicate, while error bars below the bars indicate the spread of high-confidence peptide detections with intensity coefficient of variances below 20%.

## CONCLUSIONS

LIT mass analyzers are user-friendly for maintenance and more economical to run due to less stringent vacuum requirements than OT-based mass spectrometers. Particularly in the context of single-cell proteomics where thousands of injections are required for each biological sample, the greater stability and lower expense of a LIT-based mass spectrometer make it a platform worth considering. Here we optimized acquisition methods to take advantage of a potential standalone LIT mass spectrometer for low-input proteomics and assessed those methods based on detection and quantitation accuracy. We demonstrated multiple low-input library generation schemes and show that they can be used to improve detections in single-cell experiments. Based on these results, we believe that LIT instrument platforms could give more laboratories democratized access to perform low-input proteomics analysis without requiring high-resolution mass spectrometers.

## ASSOCIATED CONTENT

### Data availability

All mass spectrometry raw data and search engine files from Spectronaut from this study have been deposited to the ProteomeXchange Consortium via the MassIVE repository. Project accession: PXD038086.

### Supporting information

Supporting table S1: Detected peptides from different scanning modes on the MS1 and MS2 levels from 1, 10, and 100 ng injection material of HeLa protein digest; Acquisition parameters for low-input DIA analysis.

## AUTHOR INFORMATION

### Authors

**Teeradon Phlairaharn** − The Novo Nordisk Foundation Center for Protein Research, Faculty of Health Sciences, University of Copenhagen, Copenhagen, Denmark; Technical University of Munich, Germany; TUM School of Natural Sciences, Department of Bioscience; Computational Medicine, Berlin Institute of Health at Charité–Universitätsmedizin Berlin, Berlin, Germany; Department of Biomedical Informatics, The Ohio State University, Columbus, Ohio 43210, United States;

**Zilu Ye** − The Novo Nordisk Foundation Center for Protein Research, Faculty of Health Sciences, University of Copenhagen, Copenhagen, Denmark;

**Elena Krismer** − The Novo Nordisk Foundation Center for Protein Research, Faculty of Health Sciences, University of Copenhagen, Copenhagen, Denmark;

**Anna-Kathrine Pedersen**− The Novo Nordisk Foundation Center for Protein Research, Faculty of Health Sciences, University of Copenhagen, Copenhagen, Denmark;

**Maik Pietzner** − Berlin Institute of Health at Charité–Universitätsmedizin Berlin, Berlin, Germany;

### Author Contributions

T.P. and B.C.S. designed the study. T.P., Z.Y., A.K.P, and E.M.S. performed the experiments. T.P., E.K., and B.C.S. performed the data analysis. T.P., Z.Y., E.K., M.P., J.V.O., E.M.S., and B.C.S. contributed with input to the method design and data evaluation. T.P., Z.Y., J.V.O., E.M.S., and B.C.S. drafted and revised the manuscript, which has been read and approved by all authors. B.C.S., E.M.S., and J.V.O. supervised the work.

## ACKNOWLEDGMENTS

T.P. would like to thank J.A. Madsen at The Novo Nordisk Foundation Center for Protein Research and L.R. Woltereck at the Technical University of Munich for technical support and R. Bruderer and O. Bernhardt from Biognosys AG, P. E. Geyer at the Exact Sciences and K. B. Emdal at The Novo Nordisk Foundation Center for Protein Research for fruitful discussions. E.K. would like to thank M. Mann for financial support, M. Strauss for fruitful discussions, and S. Grégoire for his input on data visualization.

## ABBREVIATIONS

DDA: data-dependent acquisition
DIA: data-independent acquisition
GPF: gas-phase fractionation
IT: injection time
LC: liquid chromatography
LIT: linear ion trap
MS: mass spectrometry
OHPF: off-line moderately high-pH reversed-phase fractionation
OT: orbitrap
SCP-MS: MS-based single-cell proteomics
SPD: samples per day

## REFERENCES

(1) Bache, N.; Geyer, P. E.; Bekker-Jensen, D. B.; Hoerning, O.; Falkenby, L.; Treit, P. V.; Doll, S.; Paron, I.; Müller, J. B.; Meier, F.; Olsen, J. V.; Vorm, O.; Mann, M. A Novel LC System Embeds Analytes in Pre-Formed Gradients for Rapid, Ultra-Robust Proteomics *. Mol. Cell. Proteomics 2018, 17 (11), 2284–2296.

(2) Brunner, A.-D.; Thielert, M.; Vasilopoulou, C.; Ammar, C.; Coscia, F.; Mund, A.; Hoerning, O. B.; Bache, N.; Apalategui, A.; Lubeck, M.; Richter, S.; Fischer, D. S.; Raether, O.; Park, M. A.; Meier, F.; Theis, F. J.; Mann, M.Ultra-High Sensitivity Mass Spectrometry Quantifies Single-Cell Proteome Changes upon Perturbation. Mol. Syst. Biol. 2022, 18 (3), e10798.

(3) Goebel-Stengel, M.; Stengel, A.; Taché, Y.; Reeve, J. R., Jr. The Importance of Using the Optimal Plasticware and Glassware in Studies Involving Peptides. Anal. Biochem. 2011, 414 (1), 38–46.

(4) Zhu, Y.; Piehowski, P. D.; Zhao, R.; Chen, J.; Shen, Y.; Moore, R. J.; Shukla, A. K.; Petyuk, V. A.; Campbell-Thompson, M.; Mathews, C. E.; Smith, R. D.; Qian, W.-J.; Kelly, R. T. Nanodroplet Processing Platform for Deep and Quantitative Proteome Profiling of 10–100 Mammalian Cells. Nat. Commun. 2018, 9 (1), 1–10.

(5) Woo, J.; Williams, S. M.; Markillie, L. M.; Feng, S.; Tsai, C.-F.; Aguilera-Vazquez, V.; Sontag, R. L.; Moore, R. J.; Hu, D.; Mehta, H. S.; Cantlon-Bruce, J.; Liu, T.; Adkins, J. N.; Smith, R. D.; Clair, G. C.; Pasa-Tolic, L.; Zhu, Y. High-Throughput and High-Efficiency Sample Preparation for Single-Cell Proteomics Using a Nested Nanowell Chip. Nat. Commun. 2021, 12 (1), 6246.

(6) Petrosius, V.; Aragon-Fernandez, P.; Üresin, N.; Phlairaharn, T.; Furtwängler, B.; op de Beeck, J.; Thomsen, S. F.; Keller, U. auf D.; Porse, B. T.; Schoof, E. M. Enhancing Single-Cell Proteomics through Tailored Data-Independent Acquisition and Micropillar Array-Based Chromatography. bioRxiv, 2022, 2022.11.29.518366. https://doi.org/10.1101/2022.11.29.518366.

(7) Budnik, B.; Levy, E.; Harmange, G.; Slavov, N. SCoPE-MS: Mass Spectrometry of Single Mammalian Cells Quantifies Proteome Heterogeneity during Cell Differentiation. Genome Biol. 2018, 19 (1), 161.

(8) Schoof, E. M.; Furtwängler, B.; Üresin, N.; Rapin, N.; Savickas, S.; Gentil, C.; Lechman, E.; Keller, U. A. D.; Dick, J. E.; Porse, B. T. Quantitative Single-Cell Proteomics as a Tool to Characterize Cellular Hierarchies. Nat. Commun. 2021, 12 (1), 3341.

(9) Petelski, A. A.; Emmott, E.; Leduc, A.; Huffman, R. G.; Specht, H.; Perlman, D. H.; Slavov, N. Multiplexed Single-Cell Proteomics Using SCoPE2. Nat. Protoc. 2021, 16 (12), 5398–5425.

(10) Derks, J.; Leduc, A.; Wallmann, G.; Huffman, R. G.; Willetts, M.; Khan, S.; Specht, H.; Ralser, M.; Demichev, V.; Slavov, N. Increasing the Throughput of Sensitive Proteomics by plexDIA. Nat. Biotechnol. 2022. https://doi.org/10.1038/s41587-022-01389-w.

(11) Stejskal, K.; Op de Beeck, J.; Dürnberger, G.; Jacobs, P.; Mechtler, K. Ultrasensitive NanoLC-MS of Subnanogram Protein Samples Using Second Generation Micropillar Array LC Technology with Orbitrap Exploris 480 and FAIMS PRO. Anal. Chem. 2021, 93 (25), 8704–8710.

(12) Slavov, N. Scaling Up Single-Cell Proteomics. Mol. Cell. Proteomics 2022, 21 (1), 100179.

(13) Cheung, T. K.; Lee, C.-Y.; Bayer, F. P.; McCoy, A.; Kuster, B.; Rose, C. M. Defining the Carrier Proteome Limit for Single-Cell Proteomics. Nat. Methods 2021, 18 (1), 76–83.

(14) Ye, Z.; Batth, T. S.; Rüther, P.; Olsen, J. V. A Deeper Look at Carrier Proteome Effects for Single-Cell Proteomics. Commun Biol 2022, 5 (1), 150.

(15) Gillet, L. C.; Navarro, P.; Tate, S.; Röst, H.; Selevsek, N.; Reiter, L.; Bonner, R.; Aebersold, R. Targeted Data Extraction of the MS/MS Spectra Generated by Data-Independent Acquisition: A New Concept for Consistent and Accurate Proteome Analysis. Mol. Cell. Proteomics 2012, 11 (6), pO111.016717.

(16) Ludwig, C.; Gillet, L.; Rosenberger, G.; Amon, S.; Collins, B. C.; Aebersold, R. Data-Independent Acquisition-Based SWATH-MS for Quantitative Proteomics: A Tutorial. Mol. Syst. Biol. 2018, 14 (8), e8126.

(17) Saha-Shah, A.; Esmaeili, M.; Sidoli, S.; Hwang, H.; Yang, J.; Klein, P. S.; Garcia, B. A. Single Cell Proteomics by Data-Independent Acquisition To Study Embryonic Asymmetry in Xenopus Laevis. Anal. Chem. 2019, 91 (14), 8891–8899.

(18) Venable, J. D.; Dong, M.-Q.; Wohlschlegel, J.; Dillin, A.; Yates, J. R. Automated Approach for Quantitative Analysis of Complex Peptide Mixtures from Tandem Mass Spectra. Nat. Methods 2004, 1 (1), 39–45.

(19) Douglas, D. J.; Frank, A. J.; Mao, D. Linear Ion Traps in Mass Spectrometry. Mass Spectrom. Rev. 2005, 24 (1), 1–29.

(20) Olsen, J. V.; Schwartz, J. C.; Griep-Raming, J.; Nielsen, M. L.; Damoc, E.; Denisov, E.; Lange, O.; Remes, P.; Taylor, D.; Splendore, M.; Wouters, E. R.; Senko, M.; Makarov, A.; Mann, M.; Horning, S. A Dual Pressure Linear Ion Trap Orbitrap Instrument with Very High Sequencing Speed*. Mol. Cell. Proteomics 2009, 8 (12), 2759–2769.

(21) Hebert, A. S.; Merrill, A. E.; Bailey, D. J.; Still, A. J.; Westphall, M. S.; Strieter, E. R.; Pagliarini, D. J.; Coon, J. J. Neutron-Encoded Mass Signatures for Multiplexed Proteome Quantification. Nat. Methods 2013, 10 (4), 332–334.

(22) Liu, J. M.; Sweredoski, M. J.; Hess, S. Improved 6-Plex Tandem Mass Tags Quantification Throughput Using a Linear Ion Trap-High-Energy Collision Induced Dissociation MS(3) Scan. Anal. Chem. 2016, 88 (15), 7471–7475.

(23) Park, J.; Yu, F.; Fulcher, J. M.; Williams, S. M.; Engbrecht, K.; Moore, R. J.; Clair, G. C.; Petyuk, V.; Nesvizhskii, A. I.; Zhu, Y. Evaluating Linear Ion Trap for MS3-Based Multiplexed Single-Cell Proteomics. Anal. Chem. 2023.https://doi.org/10.1021/acs.analchem.2c03739.

(24) Heil, L. R.; Remes,P. M.; MacCoss, M. J. Comparison of Unit Resolution Versus High-Resolution Accurate Mass for Parallel Reaction Monitoring. J. Proteome Res. 2021, 20 (9), 4435–4442.

(25) Borràs, E.; Pastor, O.; Sabidó, E. Use of Linear Ion Traps in Data-Independent Acquisition Methods Benefits Low-Input Proteomics. Anal. Chem. 2021, 93 (34), 11649–11653.

(26) Phlairaharn, T.; Grégoire, S.; Woltereck, L. R.; Petrosius, V.; Furtwängler, B.; Searle, B. C.; Schoof, E. M. High Sensitivity Limited Material Proteomics Empowered by Data-Independent Acquisition on Linear Ion Traps. J. Proteome Res. 2022. https://doi.org/10.1021/acs.jproteome.2c00376.

(27) Tsou, C.-C.; Avtonomov, D.; Larsen, B.; Tucholska, M.; Choi, H.; Gingras, A.-C.; Nesvizhskii, A. I. DIA-Umpire: Comprehensive Computational Framework for Data-Independent Acquisition Proteomics. Nat. Methods 2015, 12 (3), 258–264.

(28) Ting, Y. S.; Egertson, J. D.; Payne, S. H.; Kim, S.; MacLean, B.; Käll, L.; Aebersold, R.; Smith, R. D.; Noble, W. S.; MacCoss, M. J. Peptide-Centric Proteome Analysis: An Alternative Strategy for the Analysis of Tandem Mass Spectrometry Data. Mol. Cell. Proteomics 2015, 14 (9), 2301–2307.

(29) Hao, P.; Guo, T.; Li, X.; Adav, S. S.; Yang, J.; Wei, M.; Sze, S. K. Novel Application of Electrostatic Repulsion-Hydrophilic Interaction Chromatography (ERLIC) in Shotgun Proteomics: Comprehensive Profiling of Rat Kidney Proteome. J. Proteome Res. 2010, 9 (7), 3520–3526.

(30) Di Palma, S.; Boersema, P. J.; Heck, A. J. R.; Mohammed, S. Zwitterionic Hydrophilic Interaction Liquid Chromatography (ZIC-HILIC and ZIC-cHILIC) Provide High Resolution Separation and Increase Sensitivity in Proteome Analysis. Anal. Chem. 2011, 83 (9), 3440–3447.

(31) Batth, T. S.; Francavilla, C.; Olsen, J. V. Off-Line High-pH Reversed-Phase Fractionation for in-Depth Phosphoproteomics. J. Proteome Res. 2014, 13 (12), 6176–6186.

(32) Kulak, N. A.; Geyer, P. E.; Mann, M. Loss-Less Nano-Fractionator for High Sensitivity, High Coverage Proteomics*. Mol. Cell. Proteomics 2017, 16 (4), 694–705.

(33) Deutsch, E. W.; Perez-Riverol, Y.; Chalkley, R. J.; Wilhelm, M.; Tate, S.; Sachsenberg, T.; Walzer, M.; Käll, L.; Delanghe, B.; Böcker, S.; Schymanski, E. L.; Wilmes, P.; Dorfer, V.; Kuster, B.; Volders, P.-J.; Jehmlich, N.; Vissers, J. P. C.; Wolan, D. W.; Wang, A. Y.; Mendoza, L.; Shofstahl, J.; Dowsey, A. W.; Griss, J.; Salek, R. M.; Neumann, S.; Binz, P.-A.; Lam, H.; Vizcaíno, J. A.; Bandeira, N.; Röst, H. Expanding the Use of Spectral Libraries in Proteomics. J. Proteome Res. 2018, 17 (12), 4051–4060.

(34) Ezzoukhry, Z.; Henriet, E.; Cordelières, F. P.; Dupuy, J.-W.; Maître, M.; Gay, N.; Di-Tommaso, S.; Mercier, L.; Goetz, J. G.; Peter, M.; Bard, F.; Moreau, V.; Raymond, A.-A.; Saltel, F. Combining Laser Capture Microdissection and Proteomics Reveals an Active Translation Machinery Controlling Invadosome Formation. Nat. Commun. 2018, 9 (1), 1–11.

(35) Zhu, Y.; Dou, M.; Piehowski, P. D.; Liang, Y.; Wang, F.; Chu, R. K.; Chrisler, W. B.; Smith, J. N.; Schwarz, K. C.; Shen, Y.; Shukla, A. K.; Moore, R. J.; Smith, R. D.; Qian, W.-J.; Kelly, R. T. Spatially Resolved Proteome Mapping of Laser Capture Microdissected Tissue with Automated Sample Transfer to Nanodroplets*. Mol. Cell. Proteomics 2018, 17 (9), 1864–1874.

(36) Kim, Y. J.; Sweet, S. M. M.; Egertson, J. D.; Sedgewick, A. J.; Woo, S.; Liao, W.-L.; Merrihew, G. E.; Searle, B. C.; Vaske, C.; Heaton, R.; MacCoss, M. J.; Hembrough, T. Data-Independent Acquisition Mass Spectrometry To Quantify Protein Levels in FFPE Tumor Biopsies for Molecular Diagnostics. J. Proteome Res. 2019, 18 (1), 426–435.

(37) Amon, S.; Meier-Abt, F.; Gillet, L. C.; Dimitrieva, S.; Theocharides, A. P. A.; Manz, M. G.; Aebersold, R. Sensitive Quantitative Proteomics of Human Hematopoietic Stem and Progenitor Cells by Data-Independent Acquisition Mass Spectrometry. Mol. Cell. Proteomics 2019, 18 (7), 1454–1467.

(38) Spahr, C. S.; Davis, M. T.; McGinley, M. D.; Robinson, J. H.; Bures, E. J.; Beierle, J.; Mort, J.; Courchesne, P. L.; Chen, K.; Wahl, R. C.; Yu, W.; Luethy, R.; Patterson, S. D. Towards Defining the Urinary Proteome Using Liquid Chromatography-Tandem Mass Spectrometry. I. Profiling an Unfractionated Tryptic Digest. Proteomics 2001, 1 (1), 93–107.

(39) Panchaud, A.; Scherl, A.; Shaffer, S. A.; von Haller, P. D.; Kulasekara, H. D.; Miller, S. I.; Goodlett, D. R. Precursor Acquisition Independent from Ion Count: How to Dive Deeper into the Proteomics Ocean. Anal. Chem. 2009, 81 (15), 6481–6488.

(40) Ting, Y. S.; Egertson, J. D.; Bollinger, J. G.; Searle, B. C.; Payne, S. H.; MacCoss, M. J. PECAN: Library-Free Peptide Detection for Data-Independent Acquisition Tandem Mass Spectrometry Data. Nat. Methods 2017, 14 (9), 903–908.

(41) Boekweg, H.; Van Der Watt, D.; Truong, T.; Johnston, S. M.; Guise, A. J.; Plowey, E. D.; Kelly, R. T.; Payne, S. H. Features of Peptide Fragmentation Spectra in Single-Cell Proteomics. J. Proteome Res. 2022, 21 (1), 182–188.

(42) Bruderer, R.; Bernhardt, O. M.; Gandhi, T.; Miladinović, S. M.; Cheng, L.-Y.; Messner, S.; Ehrenberger, T.; Zanotelli, V.; Butscheid, Y.; Escher, C.; Vitek, O.; Rinner, O.; Reiter, L. Extending the Limits of Quantitative Proteome Profiling with Data-Independent Acquisition and Application to Acetaminophen-Treated Three-Dimensional Liver Microtissues. Mol. Cell. Proteomics 2015, 14 (5), 1400–1410.

(43) Cox, J.; Mann, M. MaxQuant Enables High Peptide Identification Rates, Individualized P.p.b.-Range Mass Accuracies and Proteome-Wide Protein Quantification. Nat. Biotechnol. 2008, 26 (12), 1367–1372.

(44) Searle, B. C.; Pino, L. K.; Egertson, J. D.; Ting, Y. S.; Lawrence, R. T.; MacLean, B. X.; Villén, J.; MacCoss, M. J. Chromatogram Libraries Improve Peptide Detection and Quantification by Data Independent Acquisition Mass Spectrometry. Nat. Commun. 2018, 9 (1), 1–12.

(45) Krieger, J. R.; Wybenga-Groot, L. E.; Tong, J.; Bache, N.; Tsao, M. S.; Moran, M. F. Evosep One Enables Robust Deep Proteome Coverage Using Tandem Mass Tags While Significantly Reducing Instrument Time. J. Proteome Res. 2019, 18 (5), 2346–2353.

(46) Granholm, V.; Noble, W. S.; Käll, L. On Using Samples of Known Protein Content to Assess the Statistical Calibration of Scores Assigned to Peptide-Spectrum Matches in Shotgun Proteomics. J. Proteome Res. 2011, 10 (5), 2671–2678.

(47) Granholm, V.; Navarro, J. F.; Noble, W. S.; Käll, L. Determining the Calibration of Confidence Estimation Procedures for Unique Peptides in Shotgun Proteomics. J. Proteomics 2013, 80, 123–131.

(48) The, M.; MacCoss, M. J.; Noble, W. S.; Käll, L. Fast and Accurate Protein False Discovery Rates on Large-Scale Proteomics Data Sets with Percolator 3.0. J. Am. Soc. Mass Spectrom. 2016, 27 (11), 1719– 1727.

(49) Feng, X.-D.; Li, L.-W.; Zhang, J.-H.; Zhu, Y.-P.; Chang, C.; Shu, K.-X.; Ma, J. Using the Entrapment Sequence Method as a Standard to Evaluate Key Steps of Proteomics Data Analysis Process. BMC Genomics 2017, 18 (Suppl 2), 143.

(50) Lin, A.; Short, T.; Noble, W. S.; Keich, U. Improving Peptide-Level Mass Spectrometry Analysis via Double Competition. J. Proteome Res. 2022, 21 (10), 2412–2420.

(51) The, M.; Samaras, P.; Kuster, B.; Wilhelm, M. Re-Analysis of ProteomicsDB Using an Accurate, Sensitive and Scalable False Discovery Rate Estimation Approach for Protein Groups. Mol. Cell. Proteomics 2022, 100437.

(52) Elias, J. E.; Gygi, S. P. Target-Decoy Search Strategy for Increased Confidence in Large-Scale Protein Identifications by Mass Spectrometry. Nat. Methods 2007, 4 (3), 207–214.

(53) Pino, L. K.; Just, S. C.; MacCoss, M. J.; Searle, B. C. Acquiring and Analyzing Data Independent Acquisition Proteomics Experiments without Spectrum Libraries. Mol. Cell. Proteomics 2020. https://doi.org/10.1074/mcp.P119.001913.

(54) Pino, L. K.; Searle, B. C.; Yang, H.-Y.; Hoofnagle, A. N.; Noble, W. S.; MacCoss, M. J. Matrix-Matched Calibration Curves for Assessing Analytical Figures of Merit in Quantitative Proteomics. J. Proteome Res. 2020, 19 (3), 1147–1153.

(55) Penny, J.; Schroeder, G. N.; Bengoechea, J. A.; Collins, B. C. A Gas Phase Fractionation Acquisition Scheme Integrating Ion Mobility for Rapid diaPASEF Library Generation. bioRxiv, 2022, 2022.07.21.500948. https://doi.org/10.1101/2022.07.21.500948.

(56) Kirkland, J. J.; van Straten, M. A.; Claessens, H. A. High pH Mobile Phase Effects on Silica-Based Reversed-Phase High-Performance Liquid Chromatographic Columns. J. Chromatogr. A 1995, 691 (1), 3–19.

(57) Pellett, J.; Lukulay, P.; Mao, Y.; Bowen, W.; Reed, R.; Ma, M.; Munger, R. C.; Dolan, J. W.; Wrisley, L.; Medwid, K.; Toltl, N. P.; Chan, C. C.; Skibic, M.; Biswas, K.; Wells, K. A.; Snyder, L. R. “Orthogonal” Separations for Reversed-Phase Liquid Chromatography. J. Chromatogr. A 2006, 1101 1-2), 122–135.

(58) Leduc, A.; Gray Huffman, R.; Cantlon, J.; Khan, S.; Slavov, N. Exploring Functional Protein Covariation across Single Cells Using nPOP. bioRxiv, 2022, 2021.04.24.441211. https://doi.org/10.1101/2021.04.24.441211.

(59) Siyal, A. A.; Chen, E. S.-W.; Chan, H.-J.; Kitata, R. B.; Yang, J.-C.; Tu, H.-L.; Chen, Y.-J. Sample Size-Comparable Spectral Library Enhances Data-Independent Acquisition-Based Proteome Coverage of Low-Input Cells. Anal. Chem. 2021, 93 (51), 17003–17011.

