## Supplementary Information for "Optimizing linear ion trap data independent acquisition towards single cell proteomics"

#### Supporting tables

Table S1

**Supporting table S1 Identified peptides from different scanning modes on the MS1 and MS2 levels from 1-, 10-, and 100 ng injection material of HeLa protein digest.**

| Input | MS1 scanning mode | Reached max fill time | Number of MS1 scans | MS2 scanning mode | Reached max fill time | Number of MS2 scans | WIndow size | Cycle time | Points per peak | Identified peptides | Identified proteins |
|---|---|---|---|---|---|---|---|---|---|---|---|
| **1 ng** | ***Zoom*** | **28.4 %** | **655** | ***Zoom*** | **16.8 %** | **3,276** | **80 m/z** | **2.8 s** | **10** | **979** | **437** |
| **1 ng** | ***Zoom*** | **40 %** | **762** | ***Enhanced*** | **82.9 %** | **12,202** | **25 m/z** | **2.4 s** | **9** | **2,502** | **995** |
| **1 ng** | ***Zoom*** | **49.3 %** | **809** | ***Normal*** | **99.8 %** | **32,360** | **10 m/z** | **2.3 s** | **9** | **3,217** | **1,158** |
| **1 ng** | ***Zoom*** | **66.3 %** | **827** | ***Rapid*** | **100 %** | **47,966** | **7 m/z** | **2.3 s** | **9** | **3,238** | **1,163** |
| **1 ng** | ***Zoom*** | **77,1 %** | **782** | ***Turbo*** | **100 %** | **62,586** | **5 m/z** | **2.3 s** | **7** | **2,736** | **1,049** |
| **1 ng** | ***Enhanced*** | **59.5 %** | **892** | ***Normal*** | **99.9 %** | **35,706** | **10 m/z** | **2 s** | **10** | **3,441** | **1,302** |
| **1 ng** | ***Normal*** | **100 %** | **910** | ***Normal*** | **100 %** | **36,400** | **10 m/z** | **2 s** | **8** | **2,499** | **909** |
| **10 ng** | ***Zoom*** | **26 %** | **785** | ***Turbo*** | **99.7 %** | **62,800** | **5 m/z** | **2.3 s** | **8** | **6,304** | **1,921** |
| **10 ng** | ***Turbo*** | **25.5 %** | **830** | ***Rapid*** | **99.1 %** | **48,140** | **7 m/z** | **2.2 s** | **9** | **6,296** | **1,866** |
| **10 ng** | ***Turbo*** | **26 %** | **817** | ***Normal*** | **87.2 %** | **32,693** | **10 m/z** | **2.2 s** | **9** | **5,270** | **1,796** |
| **10 ng** | ***Zoom*** | **24.6 %** | **783** | ***Enhanced*** | **55.4 %** | **12,533** | **25 m/z** | **2.3 s** | **9** | **3,255** | **1,226** |

| | | | | | | | | | | | |
|---|---|---|---|---|---|---|---|---|---|---|---|
| **10 ng** | ***Zoom*** | **23.2 %** | **659** | ***Zoom*** | **16.3 %** | **3,295** | **80 m/z** | **2.8 s** | **9** | **1,168** | **534** |
| **10 ng** | ***Normal*** | **34.3 %** | **937** | ***Rapid*** | **99.9 %** | **54,384** | **7 m/z** | **1.9 s** | **8** | **5,157** | **1,722** |
| **100 ng** | ***Rapid*** | **21.3 %** | **954** | ***Rapid*** | **88.4 %** | **55,332** | **7 m/z** | **1.9 s** | **10** | **6,934** | **2,340** |
| **100 ng** | ***Rapid*** | **21.8 %** | **887** | ***Turbo*** | **99 %** | **70,986** | **5 m/z** | **2.0 s** | **8** | **5,866** | **2,094** |

### Supporting methods

#### Acquisition parameters for low-input DIA analysis

##### DIA-LIT (MS1: *Zoom*, MS2: *Zoom*)

The scan sequence began with an MS1 spectrum on LIT with *Zoom* scanning speed (Scan range 400–1000 Th, automatic gain control (AGC) target of 300%, maximum injection time 10 ms, RF lens 40%). The precursor's mass range for MS2 analysis on LIT with *Zoom* scanning speed was set from 500 to 900 Th and the scan range from 200 - 1200 Th. MS2 analysis consisted of higher-energy collisional dissociation (HCD), and MS2 AGC was set to standard. the isolation window was set to 80 m/z.

##### DIA-LIT (MS1: *Zoom*, MS2: *Enhanced*)

The scan sequence began with an MS1 spectrum on LIT with *Zoom* scanning speed (Scan range 400–1000 Th, automatic gain control (AGC) target of 300%, maximum injection time 10 ms, RF lens 40%). The precursor's mass range for MS2 analysis on LIT with *Enhanced* scanning speed was set from 500 to 900 Th and the scan range from 200 - 1200 Th. MS2 analysis consisted of higher-energy collisional dissociation (HCD), and MS2 AGC was set to standard. the isolation window was set to 25 m/z.

##### DIA-LIT (MS1: *Zoom*, MS2: *Normal*)

The scan sequence began with an MS1 spectrum on LIT with *Zoom* scanning speed (Scan range 400–1000 Th, automatic gain control (AGC) target of 300%, maximum injection time 10 ms, RF lens 40%). The precursor's mass range for MS2 analysis on LIT with *Normal* scanning speed was set from 500 to 900 Th and the scan range from 200 - 1200 Th. MS2 analysis consisted of higher-energy collisional dissociation (HCD), and MS2 AGC was set to standard. the isolation window was set to 10 m/z.

##### DIA-LIT (MS1: *Zoom*, MS2: *Rapid*)

The scan sequence began with an MS1 spectrum on LIT with *Zoom* scanning speed (Scan range 400–1000 Th, automatic gain control (AGC) target of 300%, maximum injection time 10 ms, RF lens 40%). The precursor's mass range for MS2 analysis on LIT with *Rapid* scanning speed was set from 500 to 900 Th and the scan range from 200 - 1200 Th. MS2 analysis consisted of higher-energy collisional dissociation (HCD), and MS2 AGC was set to standard. the isolation window was set to 7 m/z.

DIA-LIT (MS1: *Zoom*, MS2: *Turbo*)

The scan sequence began with an MS1 spectrum on LIT with *Zoom* scanning speed (Scan range 400–1000 Th, automatic gain control (AGC) target of 300%, maximum injection time 10 ms, RF lens 40%). The precursor's mass range for MS2 analysis on LIT with *Turbo* scanning speed was set from 500 to 900 Th and the scan range from 200 - 1200 Th. MS2 analysis consisted of higher-energy collisional dissociation (HCD), and MS2 AGC was set to standard. the isolation window was set to 5 m/z.

DIA-LIT (MS1: *Enhanced*, MS2: *Normal*)

The scan sequence began with an MS1 spectrum on LIT with *Enhanced* scanning speed (Scan range 400–1000 Th, automatic gain control (AGC) target of 300%, maximum injection time 10 ms, RF lens 40%). The precursor's mass range for MS2 analysis on LIT with *Normal* scanning speed was set from 500 to 900 Th and the scan range from 200 - 1200 Th. MS2 analysis consisted of higher-energy collisional dissociation (HCD), and MS2 AGC was set to standard. the isolation window was set to 10 m/z.

DIA-LIT (MS1: *Normal*, MS2: *Normal*)

The scan sequence began with an MS1 spectrum on LIT with *Normal* scanning speed (Scan range 400–1000 Th, automatic gain control (AGC) target of 300%, maximum injection time 10 ms, RF lens 40%). The precursor's mass range for MS2 analysis on LIT with *Normal* scanning speed was set from 500 to 900 Th and the scan range from 200 - 1200 Th. MS2 analysis consisted of higher-energy collisional dissociation (HCD), and MS2 AGC was set to standard. the isolation window was set to 10 m/z.

DIA-LIT (MS1: *Turbo*, MS2: *Rapid*)

The scan sequence began with an MS1 spectrum on LIT with *Turbo* scanning speed (Scan range 400–1000 Th, automatic gain control (AGC) target of 300%, maximum injection time 10 ms, RF lens 40%). The precursor's mass range for MS2 analysis on LIT with *Rapid* scanning speed was set from 500 to 900 Th and the scan range from 200 - 1200 Th. MS2 analysis consisted of higher-energy collisional dissociation (HCD), and MS2 AGC was set to standard. the isolation window was set to 7 m/z.

DIA-LIT (MS1: *Turbo*, MS2: *Normal*)

The scan sequence began with an MS1 spectrum on LIT with *Turbo* scanning speed (Scan range 400–1000 Th, automatic gain control (AGC) target of 300%, maximum injection time 10 ms, RF lens 40%). The precursor's mass range for MS2 analysis on LIT with *Normal* scanning speed was set from 500 to 900

Th and the scan range from 200 - 1200 Th. MS2 analysis consisted of higher-energy collisional dissociation (HCD), and MS2 AGC was set to standard. the isolation window was set to 10 m/z.

DIA-LIT (MS1: *Normal*, MS2: *Rapid*)

The scan sequence began with an MS1 spectrum on LIT with *Normal* scanning speed (Scan range 400–1000 Th, automatic gain control (AGC) target of 300%, maximum injection time 10 ms, RF lens 40%). The precursor's mass range for MS2 analysis on LIT with *Rapid* scanning speed was set from 500 to 900 Th and the scan range from 200 - 1200 Th. MS2 analysis consisted of higher-energy collisional dissociation (HCD), and MS2 AGC was set to standard. the isolation window was set to 7 m/z.

DIA-LIT (MS1: *Rapid*, MS2: *Rapid*)

The scan sequence began with an MS1 spectrum on LIT with *Rapid* scanning speed (Scan range 400–1000 Th, automatic gain control (AGC) target of 300%, maximum injection time 10 ms, RF lens 40%). The precursor's mass range for MS2 analysis on LIT with *Rapid* scanning speed was set from 500 to 900 Th and the scan range from 200 - 1200 Th. MS2 analysis consisted of higher-energy collisional dissociation (HCD), and MS2 AGC was set to standard. the isolation window was set to 7 m/z.

DIA-LIT (MS1: *Rapid*, MS2: *Turbo*)

The scan sequence began with an MS1 spectrum on LIT with *Rapid* scanning speed (Scan range 400–1000 Th, automatic gain control (AGC) target of 300%, maximum injection time 10 ms, RF lens 40%). The precursor's mass range for MS2 analysis on LIT with *Turbo* scanning speed was set from 500 to 900 Th and the scan range from 200 - 1200 Th. MS2 analysis consisted of higher-energy collisional dissociation (HCD), and MS2 AGC was set to standard. the isolation window was set to 5 m/z.
